# A histone methyltransferase inhibitor can reverse epigenetically acquired drug resistance in the malaria parasite *Plasmodium falciparum*

**DOI:** 10.1101/775734

**Authors:** Amanda Chan, Alexis Dziedziech, Laura A Kirkman, Kirk W Deitsch, Johan Ankarklev

## Abstract

Malaria parasites invade and replicate within red blood cells (RBCs), extensively modifying their structure and gaining access to the extracellular environment by placing the plasmodial surface anion channel (PSAC) into the RBC membrane. Expression of members of the cytoadherence linked antigen gene 3 (*clag3*) family is required for PSAC activity, a process that is regulated epigenetically. PSAC is a well-established route of uptake for large, hydrophilic antimalarial compounds and parasites can acquire resistance by silencing *clag3* gene expression, thereby reducing drug uptake. We found that exposure to sub-IC50 concentrations of the histone methyltransferase inhibitor chaetocin caused substantial changes in both *clag3* gene expression and RBC permeability, reversing acquired resistance to the antimalarial compound blasticidin S that is transported through PSAC. Chaetocin treatment also altered progression of parasites through their replicative cycle, presumably by changing their ability to modify chromatin appropriately to enable DNA replication. These results indicate that targeting histone modifiers could represent a novel tool for reversing epigenetically acquired drug resistance in *P. falciparum*.

**Importance:** Drug resistance is a major concern for the treatment of infectious diseases throughout the world. For malaria, a novel mechanism of resistance was recently described in which epigenetic modifications led to a resistance phenotype that is rapidly reversible, thus reducing the fitness cost that is often associated with genetic mutations that lead to resistance. The possibility of this type of resistance arising in a natural setting is particularly troubling since parasites could rapidly switch to and from a resistant phenotype, thus making it especially difficult to combat. Here we show that application of a histone methyltransferase inhibitor can rapidly reverse the epigenetic changes that lead to drug resistance, thereby causing parasites to revert to a drug sensitive phenotype. This is a novel application of drugs that target epigenetic modifiers and lends additional support for ongoing efforts to develop drugs against malaria that target the histone modifiers of the parasite.

## Introduction

Malaria is a disease common in tropical and subtropical regions of the developing world where it continues to cause significant morbidity and mortality, primarily among young children. The disease is caused by infection with protozoan parasites of the genus *Plasmodium*. These parasites invade and replicate within the circulating red bloods cells (RBCs) of their hosts, causing anaemia through RBC destruction and obstructing blood flow by the adhesion of infected cells to the endothelium of post-capillary blood vessels (1). The most virulent of the human malaria parasites, *Plasmodium falciparum*, is known for making extensive modifications to the membrane of the infected RBC (2). These include changes in rigidity and flexibility (3–6), the deposition of electron-dense “knobs” within the RBC cytoskeleton (7,8), and significant changes in the permeability of the RBC membrane to various solutes, in particular anions and small compounds (9). RBC remodelling by *P. falciparum* therefore is a key aspect of parasite biology that plays a significant role in the virulence of the parasite. Targeting these modifications for novel intervention strategies is an attractive potential approach for fighting the disease. In particular, given that the increase in RBC membrane permeability provides access to the parasite, this trait could potentially be exploited in the development of new antimalarial strategies.

The changes in permeability to the RBC membrane resulting from infection by *P. falciparum* have been extensively studied. Experiments employing osmotic lysis or electrophysiology of infected cells have detected the induction by parasites of an anion pore or channel within the RBC membrane, alternatively referred to as the new permeability pathway (NPP) (10) or the plasmodial surface anion channel (PSAC) (11). A significant breakthrough in the understanding of the molecular nature of these channels came with the discovery that the parasite-encoded protein CLAG3 plays a pivotal role in RBC membrane permeability (12), and that it likely is an important component of the pore itself. Interestingly, this protein exists in two forms encoded by alternatively transcribed genes called *clag3.1* and *clag3.2*, and certain PSAC properties are determined by which gene is expressed (12,13).

Unlike the other members of the *clag* gene family, *clag3.1* and *clag3.2* display clonally variant expression and thus can exist in either a transcriptionally active or silent state, and this transcriptional state can be stably inherited through many cellular divisions. Clonally variant expression of these genes is controlled epigenetically, specifically through the incorporation of the histone mark H3K9me3 into the chromatin surrounding silent gene while the opposing mark, H3K9ac is found at transcriptionally active locus (14–16). Expression is also mutually exclusive, thus one gene displays the active H3K9ac mark while the silent modification H3K9me3 is enriched at the other allele (13,17). Switches in which gene is expressed correspond to changes in the phenotypic characteristics of PSAC. Interestingly, it was demonstrated that cultured parasites can acquire resistance to large, hydrophilic drugs like blastidiicin S or leupeptin by down-regulating PSAC activity (18,19) by simultaneously silencing both *clag3* alleles (13,20). This unusual situation resulted from the incorporation of silent histone marks at both genes, thus down-regulating pore activity and preventing drug uptake. Recent work by Mira-Martinez and colleagues further showed that antimalarial compounds known as bis-thiasolum salts require pore activity to enter infected cells, while compounds like doxycline, azithromycin and lumefantrine enter via an alternative route, thereby better defining the important role *clag3* expression plays in the acquisition of resistance to a subset compounds that kill parasites (21). The ability of parasites to control access of antimalarial compounds to the intracellular environment presents a potential new mechanism for the development of drug resistance in the field, a troubling possibility (22). However, with an understanding of the molecular basis of activation and silencing of *clag3.1/3.2*, it might be possible to manipulate *clag3.1/3.2* gene activity and thereby reverse epigenetically acquired drug resistance and potentially increase accessibility of parasites to antimalarial compounds.

Inhibitors of histone modifiers have been developed as potential therapeutic agents against numerous diseases, including malaria (23–25). The compounds typically work by altering the chromatin structure of the targeted cell sufficiently to disrupt gene expression patterns or prevent DNA replication, thus either killing the cell or preventing proliferation. Cell cycle progression in malaria parasites is unusual since unlike model eukaryotes that replicate by binary fission, malaria parasites undergo repeated rounds of unsynchronized genome replication and nuclear division in the absence of cell division (26,27), a process called schizogony. What triggers each additional round of replication and how the cycle is regulated are largely unknown, although the Plasmodium-specific kinase CRK4 was shown to play a key role in this process (28). The heterochromatin mark H3K9me3 has been shown to affect pre-initiation complex formation at origins (29) and rates of DNA replication (30) in higher eukaryotic cells, thus inhibitors that affect H3K9me3 deposition would be predicted to have interesting effects on schizogony,

In addition to their development as therapeutic compounds, inhibitors that target various aspects of chromatin structure can also be used experimentally to probe the function of specific histone modifications and decipher mechanisms that regulate patterns of gene expression or cell replication. For example, we previously utilized sub-IC50 concentrations of the histone methyltransferase inhibitor chaetocin to investigate how changes in histone methylation efficiency affected patterns of *var* gene expression switching (31). This inhibitor targets histone methyltransferases of the SET3 family that deposit the histone mark H3K9me3 (32). In *P. falciparum*, this mark is devoted almost exclusively to regulating transcription of clonally variant gene families, which in addition to *var* genes also includes *clag3.1/3.2* (15,16), thus providing a potential tool for investigating mechanisms of acquired resistance to antimalarial compounds that enter infected cells through PSAC.

Here we report that parasites grown in the presence of low doses of chaetocin display altered expression of *clag3.1/3.2*. In particular, exposure to chaetocin re-establishes PSAC activity and reverses drug resistance in parasites that have become drug-insensitive by down-regulating *clag3.1/3.2* expression. Thus, manipulation of *clag3.1/3.2* expression could represent a new avenue for delivery of antimalarial compounds to parasitized red blood cells as well as for reversing acquired drug resistance in *P. falciparum*. In addition, treatment with chaetocin slowed parasite progression through the multiple cycles of DNA replication that occur during schizogony, presumably by altering the ability of the parasites to make the appropriate chromatin modifications required for DNA replication. These observations provide insight into the poorly understood process by which parasites undergo repeated rounds of DNA replication in the absence of cell division.

## Results

### *Treatment with the histone methyltransferase inhibitor chaetocin reverses epigenetic silencing of* clag3.1/3.2 *expression*

The two alleles *clag3* are located in close proximity to one another within a ~20 kb segment of chromosome 3. The nucleosomes associated with the active or silent allele have been shown to be marked by the histone post-translational modifications H3K9ac or H3K9me3, respectively (13,17). When cultured parasites were selected for resistance to antimalarial compounds that are taken up through PSAC, both alleles were shown to acquire the H3K9me3 silencing mark, thus down-regulating total *clag3* expression, reducing pore activity and enabling the parasites to survive drug selection (13,19,20,22). Additional work showed that exposure to low doses of blasticidin, a drug known to be taken up through PSAC, selects for expression of *clag 3.1*, and silencing of both *clag3.1* and *clag3.2* when parasites are exposed to high doses of blasticidin (13). These experiments demonstrated that the two alleles differ somewhat in their ability to enable uptake of different compounds.

The small molecule chaetocin is an inhibitor of histone methyltransferases of the SET3 variety that are responsible for deposition of H3K9me3, thus treatment of cells with chaetocin can be used as a tool to investigate the role of this histone modification in controlling gene expression. Previous work showed that treatment with low levels of chaetocin induced significant changes in *var* gene expression (31), presumably by reducing incorporation of H3K9me3 into the surrounding chromatin. We were curious to see if treatment with chaetocin could similarly alter *clag3.1/3.2* expression, and more specifically, to determine if treatment with this compound could reverse the epigenetic silencing of both *clag3* alleles observed in parasites that are resistant to blasticidin. To test this hypothesis, we utilized a line of FCB parasites that had previously been selected for the development of resistance to blasticidin and shown to have repressed *clag3.1/3.2* expression (20), a kind gift from the Desai lab. We cultured blasticidin resistant parasites in the presence of sub-IC50 levels of chaetocin for two weeks, then determined steady state levels of *clag3.1* and *clag3.2* transcripts using Q-RT-PCR.

As expected, FCB wildtype parasites that had not been selected for resistance to blasticidin displayed robust *clag3* expression, with *clag3.2* being the dominantly expressed allele (Figure 1). When these parasites were selected for resistance to blasticidin, expression of *clag3.2* was dramatically reduced, leading to much lower overall *clag3* expression levels, although a low level of *clag3.1* expression remained largely unchanged. This is consistent with previous reports showing that expression of *clag3.2* is much more sensitive to blasticidin selection due to more efficient uptake of the drug when this allele is expressed (13). Removal of blasticidin pressure for 2 weeks led to a moderate increase in overall *clag3* expression as the parasites began to revert to wildtype expression levels. The dominant allele however shifted to *clag3.1*, similar to what was previously reported for *clag3* expression after removal of blasticidin selection (13). Interestingly, exposure to sub-IC50 levels of chaetocin after removal of blasticidin selection resulted in a dramatic increase in *clag3.1* expression (Figure 1), indicating that inhibition of H3K9me3 deposition reversed the epigenetic silencing of *clag3.1*, thereby reactivating the gene. It is worth noting that while overall *clag3* expression returned to levels similar to that observed in untreated parasites, expression was dominated by *clag3.1* with *clag3.2* expression levels remaining relatively low. This suggests that at this concentration of chaetocin, mutually exclusive expression remains intact and only one allele was reactivated.

**Figure 1.**
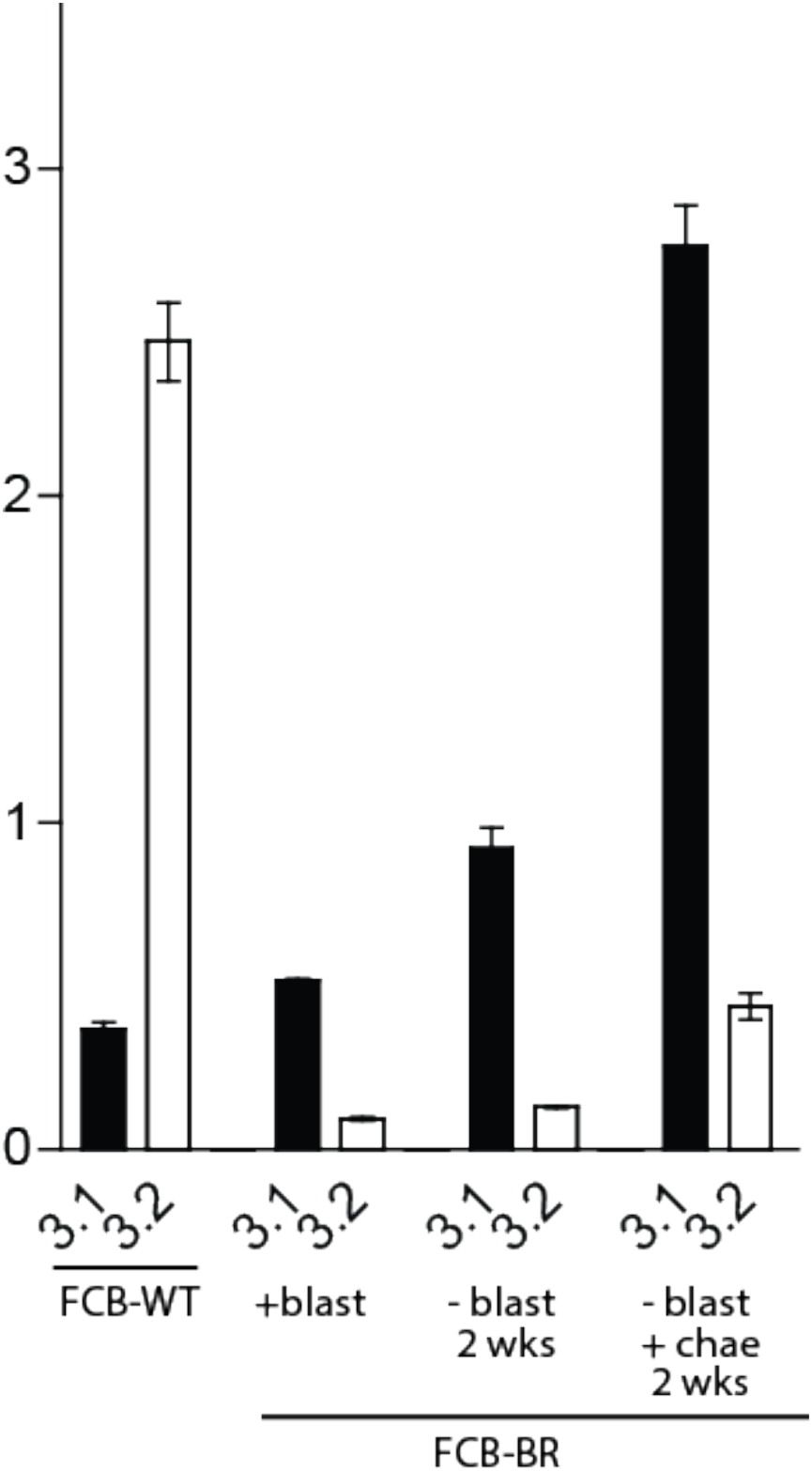
Steady state mRNA levels expressed from *clag3.1* and *clag3.2* in cultured parasites. RNA was extracted from synchronized cultures of late stage parasites from of the FCB isolate of *P. falciparum*. Levels of mRNA were determined using Q-RT-PCR from cDNA prepared from wildtype parasite (FCB-WT) or parasites that had been selected for resistance to blasticidin (FCB-BR). The resistant parasites were either cultured in the presence of blasticidin (+blast), in the absence of blasticidin for 2 weeks (-blast 2wks) or in the absence of blasticidin and in the presence of chaetocin for 2 weeks (-blast +chae 2 wks).

### Chaetocin treatment can reverse drug resistance acquired through reduced anion channel activity

In addition to blasticidin, solutes like sorbitol and alanine as well as antimalarial compounds of certain classes have also been shown to enter the infected RBC through the parasite induced anion channels, thereby gaining access to the parasite (18–22). Resistance to these compounds can be acquired by either mutations in CLAG3 that alter properties of the channel (33) or alternatively by epigenetically repressing expression of both *clag 3.1* and *clag 3.2*, as shown for the blasticidin resistant FCB line (20). The latter mechanism is particularly troubling since it could potentially lead to resistance to any antimalarial compound that enters the infected cell through the channel. Given our observations that treating parasites with sub-IC50 concentrations of chaetocin could de-repress *clag3.1/3.2* expression, we hypothesized that this could reverse resistance to drugs like blasticidin in parasites that had acquired resistance by down-regulating channel activity.

To test this hypothesis, we again utilized the blasticidin resistant line of FCB parasites that had previously been shown to have repressed *clag3.1/3.2* expression (Figure 1) (20). These parasites were grown in the presence of both chaetocin and blasticidin to determine the effect of chaetocin exposure on blasticidin resistance. The data in Figure 1 show that the effect of chaetocin on *clag3.1/3.2* expression was detectable after two weeks of exposure, however the length of time required for changes in CLAG expression and altered pore activity were not known. To determine if chaeotocin exposure alters uptake of blasticidin over time, resistant FCB parasites were cultured in the presence of either blasticidin, chaetocin or both compounds and parasite growth assayed daily by flow cytometry (Figure 2). The blasticidin resistant parasites displayed slightly slower growth in the presence of chaetocin (Figure 2A, left panel) similar to the growth observed in wildtype FCB parasites grown in the presence of chaetocin (Figure 2A, center panel). In the presence of both blasticidin and chaetocin, the FCB resistant parasites initially grew well, however after approximately 6 days of exposure to both compounds, parasite growth was arrested and the parasites failed to continue to replicate (Figure 2A, right panel), suggesting they had become sensitive to blasticidin.

**Figure 2.**
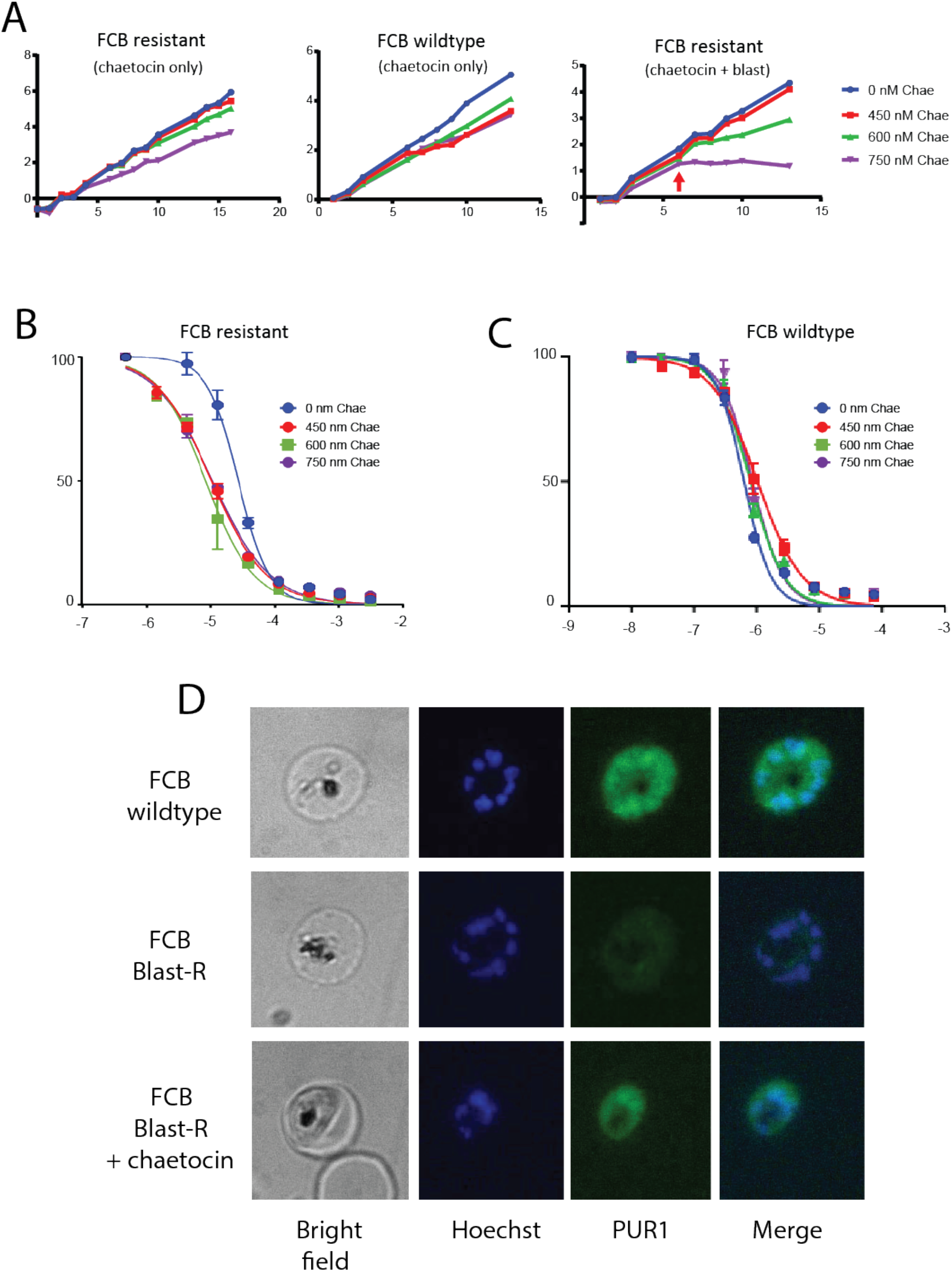
Chaetocin treatment can reverse resistance to blasticidin. Blasticidin resistant FCB parasites display reduced CLAG expression and are less sensitive to treatment with blasticidin. (A) Growth rates of resistant parasites grown in the presence of 450, 600 or 750 nM chaetocin, 2.5 μg/ml blasticidin or both compounds over time. Sensitivity to blasticidin is observed after approximately 6 days of chaetocin treatment (red arrow, right panel). (B) Sybr-green based drug sensitivity curves to determine sensitivity to blasticidin. The X-axis displays sybr green fluorescence as a percentage of that observed in parasites grown in the absence of blasticidin while the Y-axis displays the log of molar blasticidin concentration. Blasticidin resistant parasites grown in the absence of chaetocin are shown in blue while parasites grown in the presence of 450, 600 or 750 nM chaetocin for 13 days are shown in red, green or purple, respectively. (C) Blasticidin sensitivity curves for wildtype FCB parasites grown in the absence of chaetocin (blue) or in the presence of 450, 600 or 750 nM chaetocin (red, green or purple, respectively). (D) Uptake of PUR1 in wildtype (top), blasticidin resistant (middle) or blasticidin resistant parasites grown in the presence of chaetocin.

To obtain a more quantitative measurement of changes in sensitivity to blasticidin, we performed drug sensitivity assays to identify changes in IC50 resulting from chaetocin treatment. Blasticidin resistant FCB parasites were released from blasticidin pressure and were grown in the presence or absence of three different sub-IC50 concentrations of chaetocin (450, 600 and 750 nM) for thirteen days, then drug sensitivity assays were performed using a standard sybr green growth assay (34). As previously reported, this FCB line of parasites displayed a marked resistance to blasticidin, however after exposure to all three concentrations of chaetocin these parasites became much more sensitive to blasticidin and were rapidly killed by concentrations known to kill most lines of *P. falciparum* (Figure 2B). The shift in sensitivity to blasticidin resulting from chaetocin exposure was not simply the result of synergism between the two compounds since chaetocin treatment had no effect on blasticidin sensitivity of wildtype parasites (Figure 2C).

An alternative method for directly observing anion channel activity within the infected RBC membrane is via uptake of the fluorescent dye benzothiocarboxypurine, also known as PUR-1. This dye has been shown to enter infected RBCs through the parasite induced anion channel where it forms a brightly fluorescent complex with parasite nucleic acids (35). Uninfected RBCs and cells infected with ring-stage parasites do not take up the dye. To investigate changes in channel activity resulting from treatment with chaetocin, we exposed synchronized cultures of chaetocin treated and untreated parasites to PUR-1, then examined the degree of dye uptake using fluorescent microscopy. As expected, wildtype FCB parasites efficiently take up the dye and appear brightly fluorescent while blasticidin resistant parasites fail to take-up the dye and fluoresce much more weakly (Figure 2D). In contrast, blasticidin resistant parasites grown in the presence of chaetocin were brightly fluorescent, indicating that they had reactivated the pore and now readily take up PUR1 (Figure 2D). These data provide additional evidence that treatment with sub-IC50 levels of chaetocin can re-activate PSAC activity in parasites that have silenced CLAG expression.

### Chaetocin treatment alters DNA replication and transition through schizogony

Given the established role of heterochromatin and H3K9me2 on DNA replication, we examined whether treatment with sub-IC50 concentration of chaetocin might affect progression through schizogony. Parasites grown in the presence or absence of chaetocin were tightly synchronized and allowed to progress through the entire replicative cycle. Levels of DNA and RNA were assayed by flow cytometry at 2-hour intervals, thus enabling us to monitor cells as they entered and exited the asexual cycle. We observed that parasites treated with chaetocin had a reproducible delay at the onset of DNA replication, and this delay continues as the parasites progress through trophozoites and toward schizonts (Figure 3). However, by the end of the 48-hour cycle, the chaetocin-treated parasites completed replication and reinvasion at rates similar to the untreated parasites, indicating that chaetocin treatment had affected the rate of DNA replication but not the ability of the parasites to complete schizogony.

**Figure 3.**
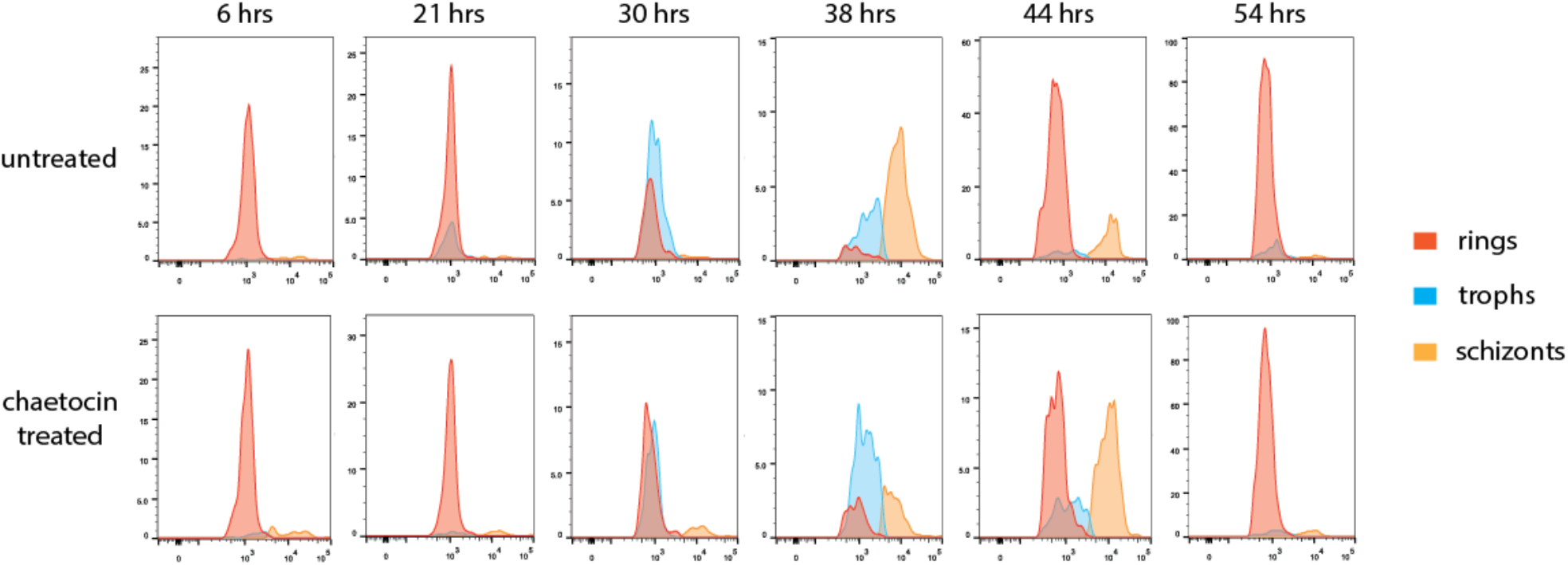
Progression through schizogony in the presence of absence of chaetocin. Cultured parasites were grown in the absence (untreated) or presence (chaetocin treated) of the histone methyltransferase inhibitor chaetocin. Cultures were tightly synchronized and monitored by flow cytometry as they progressed through the replicative cycle. Proportions of rings, trophozoites and schizonts were determined by both RNA content (thiazole orange fluorescence) and DNA content (Hoechst 33342 fluorescence) using a gating strategy described in the Methods section. The approximate time after initial red cell invasion is shown above each column.

## Discussion

The development of resistance to anti-malarial drugs remains an ongoing problem for the global effort to combat malaria. The spread of reduced sensitivity to the current most widely used antimalarial, artemisinin, have now been reported (36,37), reinforcing the notion that the development of drug resistance is an ever-present problem. The acquisition of drug resistance by altering uptake pathways through downregulation of porins has been well documented in bacterial pathogens (38), highlighting the possibility of this type of drug resistance arising for malaria parasites through the downregulation of PSAC.

Most examples of drug resistance result from genetic mutations that lead to amino acid changes in proteins that are either the direct target of the drug or that play a role in access of the drug to its target. Such genetic changes tend to be relatively stable, enabling rapid spread throughout a population but also frequently inflicting an associated fitness cost, which can lead to slow re-emergence of drug sensitivity when drug pressure is removed, as was observed for chloroquine resistance in some geographical regions (39). In contrast, the development of drug resistance through epigenetic changes, as described for the *clag3.1/3.2* locus, is potentially a more easily reversible phenotype, thus avoiding some of the fitness costs associated with heritable genetic changes. The ability of parasites to rapidly switch between resistant and sensitive phenotypes could enable them to easily adapt to the presence or absence of drug, thus further complicating malaria containment strategies. Efficient changes in gene expression patterns through epigenetic switching have been well documented for the processes of antigenic variation and sexual differentiation (40), indicating that parasites are capable of such rapid changes in gene expression. The effect of chaetocin on *clag3.1/3.2* expression however indicates that this type of epigenetic switching can itself be targeted, providing a possible method to address resistance that arises through an epigenetic mechanism.

Stanojcic et al. described in detail the unusual dynamics of DNA replication displayed by malaria parasites over the course of a cycle of schizogony (41). Surprisingly, replication velocity slowed as schizogony progressed, the opposite of what is observed during S-phase progression in mammalian cells (42). These authors offered several possible explanations for the slowing of DNA replication rates, including reduced availability of nucleotides after several rounds of replication or changes in chromatin structure within various regions of the genome. Our data indicating that chaetocin treatment slows progression through the replicative cycle is consistent with the latter hypothesis. In model eukaryotes, reduction in H3K9me3 levels, a predicted result of chaetocin treatment, increases replication rates (29,30). However, other studies have identified specific subsets of late-firing replication origins that are specifically associated with H3K9me3 (43,44), indicating that this histone mark might play a role in recruiting the replication machinery to specific regions of the genome. Future work investigating the mechanisms regulating DNA replication during schizogony by malaria parasites are likely to identify additional characteristics quite different from what would be predicted from the study of model organisms. Such differences are both interesting for the insights they provide into the evolution of cellular replication in various eukaryotic lineages as well as for the opportunities they provide for the development of novel strategies to combat malaria.

## Materials and Methods

### Parasite culture

All *P. falciparum* parasites were cultured according to standard procedures in media containing Albumax II (Gibco) without human serum. Parasites were incubated at 37°C in an atmosphere of 5% oxygen, 5% carbon dioxide, and 90% nitrogen. 3D7 parasites were obtained from MR4 (MRA-156, MR4, BEI-Resources) and blasticidin resistant FCB parasites (20) were a kind gift from Dr. Sanjay Desai at the laboratory for Malaria and Vector Research, NIAID, NIH.

### Analysis of clag3.1 and clag3.2 steady state RNA

RNA was extracted from synchronized late stage parasites 48 h after isolation of schizonts using magnetic separation (45). RNA extraction was performed using Trizol LD Reagent (Invitrogen) as described (46). RNA was purified using the PureLink RNA Mini Kit (Ambion) according to manufacturer’s protocol and afterwards treated with DNase I (Invitrogen). cDNA synthesis was performed with Superscript II RNase H reverse transcriptase (Invitrogen) with random primers (Invitrogen) as described by the manufacturer. 800 ng of total RNA were used for each cDNA synthesis reaction and a control reaction without reverse transcriptase was performed in parallel. Quantitative PCR (Q-RT-PCR) analysis was done using the relative standard curve method. All Q-RT-PCR reactions were performed in triplicate with an ABI Prism 7900HT (Applied Biosystems) using iTaq SYBR Green Supermix (Bio-Rad) and previously described primers specific for either *clag3.1* or *clag3.2* (13). Expression levels were normalized to the house keeping gene seryl tRNA synthetase.

### Uptake of PUR-1

PUR-1 uptake assays were performed as described by Kelly et al. (35). Benzothiocarbozypurine (PUR-1) was obtained from Sigma Aldrich and dissolved in ethanol at a concentration of 10 mM and diluted to 5 μM in complete media. Equal volumes of PUR-1 solution and aliquots of synchronized *P. falciparum* cultures were mixed and allowed to incubate for 10 minutes. Cells were then washed twice with culture media and observed on a Leica DMI 6000b fluorescent microscope (excitation wavelength 476 nM, emission detection between 500 to 550 nm) using a Leica DFC 360FX camera.

### Drug sensitivity assay

Drug sensitivity assays were performed as described by Smilkstein et al. (34). 100 μl aliquots of parasite culture were distributed into clear, 96 well plates at a starting parasitemia of 0.2-0.5% and 2% haematocrit. Blasticidin S was obtain from Sigma Aldrich and diluted appropriately in complete media to achieve final concentrations in a 96 well culture plate ranging from 0.01 μM to 72.9 μM for blasticidin sensitive parasites lines and 0.48 μM to 3.2 mM for blasticidin resistant parasite lines. After addition of parasites and RBCs, each well of the 96 well plate had a total volume of 200 μl per well. Plates were incubated at 37° C in an airtight chamber containing 5% oxygen, 5% carbon dioxide and 90% nitrogen. Cultures were allowed to incubate for 72 hours. To assay for growth, the cultures were resuspended and 150 μl from each well transferred to a 96 well black plate and placed at −80° C overnight. Plates were then thawed and 100 μl of SYBR green solution (0.2 μl SYBR Green/ml lysis buffer) added to each well. Plates were incubated in the dark at room temperature for one hour, shaking. SYBR Green incorporation was measured with a SpectraMax Gemini using an excitation wavelength of 490 nm and 530 nm detection. All assays were performed with triplicit wells on each plate (technical replicates) and in three independent plates (biological replicates). Data were analysed using Graphpad Prism software plotting counts against the log of the drug concentration, normalized and curve fitted by nonlinear regression (sigmoidal dose-response/variable slope equation) to yield IC50 values.

### Flow cytometry and assays for cell cycle progression

Progression through schizony was determined by flow cytometric analysis of parasite RNA & DNA content as previously described (47). Briefly, live parasite were stained at 37°C with 16 μM Hoechst 33342 and 0.1 μg/mL Thiazole Orange for 30 min at 1% hematocrit in incomplete media followed by a single wash in PBS. Cells were then diluted to 0.1% hematocrit in PBS and analyzed using a Cytek DxP11 flow cytometer for Hoechst 33342 DNA-staining (375nm laser, 450/50 emission filter) and thiazole orange RNA-staining (488nM laser, 550/30 emission filter). For each sample, 50,000 infected RBCs (DNA^+^) were gated into ring (DNA^low^/RNA^low^), trophozoite (DNA^low^/RNA^high^), and schizont stages (DNA^high^/RNA^high^).

## Acknowledgements

We thank Dr. Sanjay Desai for the kind gift of the blasticidin resistant line of FCB parasites and Dr. Bjorn Kafsack for assistance with flow cytometry and fluorescent microscopy.

